# Biases in arginine codon usage correlate with genetic disease risk

**DOI:** 10.1101/825703

**Authors:** Katharina V. Schulze, Neil A. Hanchard, Michael F. Wangler

## Abstract

**Purpose:** The persistence of hypermutable ‘CGN’ (CGG, CGA, CGC, CGU) arginine codons at high frequency suggests the possibility of negative selective pressure at these sites and that arginine codon usage could be a predictive indicator of human disease genes.

**Methods:** We analyzed arginine codons (CGN, AGG, AGA) from all ‘canonical’ Ensembl protein coding gene transcripts before comparing the frequency of CGN codons between genes with and without human disease associations and with gnomAD constraint metrics.

**Results:** The frequency of CGN codons among a gene’s total arginine codon count was higher in genes linked to syndromic autism spectrum disorder (ASD) compared to genes not associated with ASD. A comparison of genes annotated as dominant or recessive with control genes not matching either classification revealed a progressive increase in CGN codon frequency. Moreover, CGN frequency was positively correlated with a gene’s probability of loss-of-function intolerance (pLI) score and negatively correlated with ‘observed-over-expected’ ratios for both loss of function and missense mutations.

**Conclusion:** Our findings indicate that genes utilizing CGN arginine codons rather than AGG or AGA are more likely to underlie single gene disorders, particularly for dominant phenotypes, and thus constitute candidate genes for the study of human genetic disease.

## INTRODUCTION

Cytosines in a cytosine-guanine (CpG) dinucleotide context are known for their propensity to mutate at a rate that can be several hundred fold greater than transversions at other bases (*1*). This increased rate of mutation can be explained by the deamination of methylated cytosine (5mC) that results in thymine, which is not readily detected by DNA repair mechanisms (*2*). Interestingly, arginine is the only amino acid to contain CpG dinucleotides at the first and second codon positions, which are less redundant than the third position and therefore more likely to lead to an amino acid change. Moreover, codon usage for arginine can have an impact on GC content for a given gene (*3*) as arginine is encoded by six codons in vertebrate genomes – four ‘CGN’ codons (CGA, CGC, CGG, CGU) and two ‘AGR’ codons (AGG, AGA).

Based on data archived in the database of clinically relevant variants (ClinVar) (*4*), arginine substitutions underlie 20.0% of all ‘Pathogenic’ single nucleotide variants (SNVs), making arginine the most commonly substituted amino acid (**Figure 1A**). Arginine is also the most commonly substituted amino acid among ‘Benign’ variants, but at a significantly lower frequency (13.9%; Chi-squared test, p=5.53 × 10^−29^). For pathogenic as well as benign arginine substitutions, the most frequent SNVs are cytosine to thymine (C>T) and guanine to adenine (G>A) transitions – both products of 5mC deamination on the plus and minus DNA strand, respectively (**Figure 1B**).

**Figure 1.**
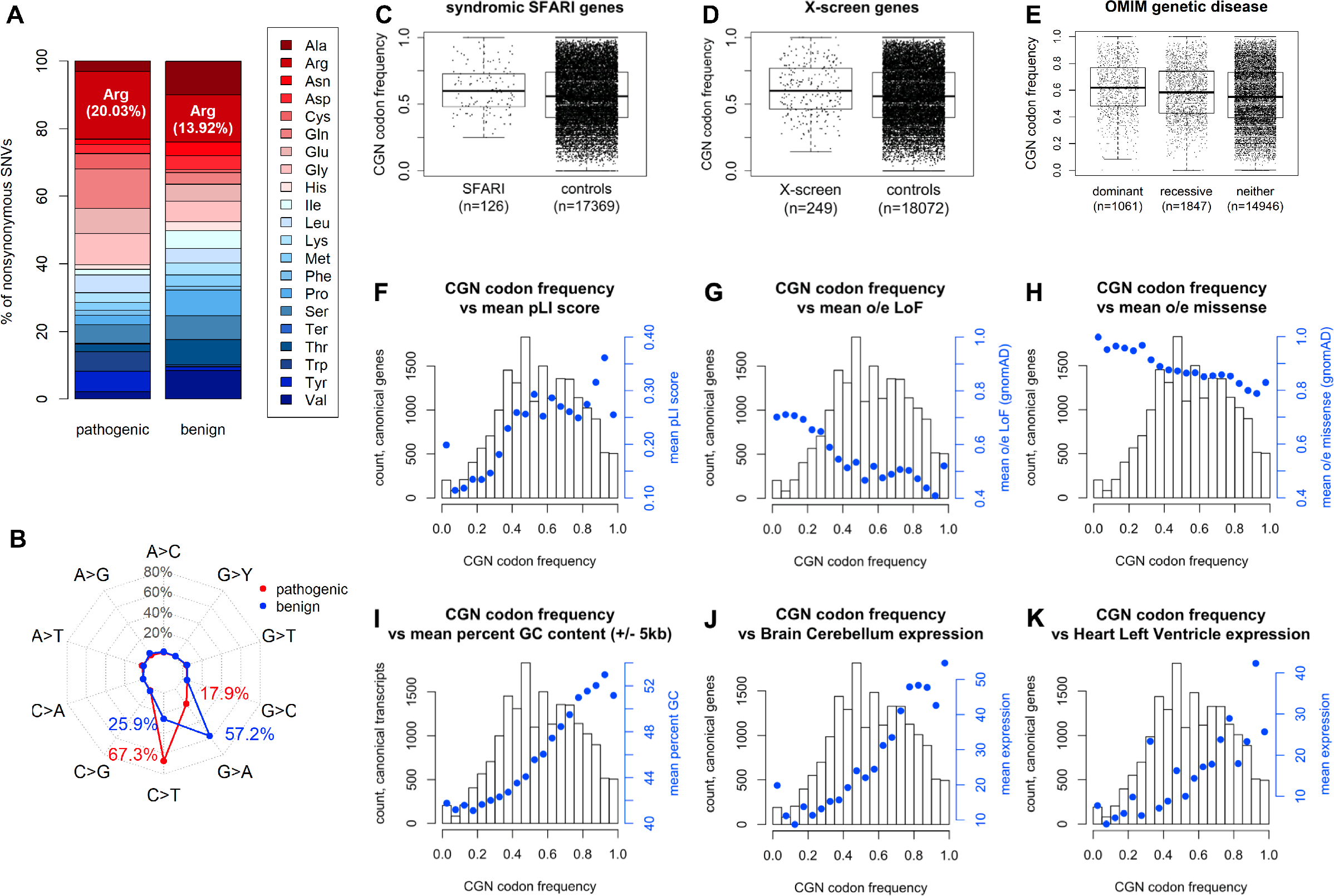
CGN arginine codon frequency and gene pathogenicity. (**A**) Amino acids with ‘Pathogenic’ and ‘Benign’ nonsynonymous single nucleotide variants (SNVs) in ClinVar. ‘Ter’ indicates a stop codon. (**B**) Arginine substituting single nucleotide changes for ‘Pathogenic’ and ‘Benign’ SNVs. (**C**) CGN arginine codon frequency comparison between syndromic SFARI genes and control genes not linked to ASD in SFARI. (**D**) CGN arginine codon frequency comparison between human homologs to essential genes detected in Drosophila X-screen and all other genes (controls). (**E**) CGN arginine codon frequency comparison between OMIM disease genes and non-disease gene controls. (**F-H**) CGN codon frequency comparison with gnomAD constraint metrics: probability of loss-of-function (pLI) score, observed over expected (o/e) ratio of loss-of-function (LoF) and o/e ratio of missense variants. (**I**) CGN codon frequency comparison with GC content 5kb up- and downstream of transcript boundaries and examples of median gene transcripts per million (TPM) for cerebellum (**J**) and left ventricle (**K**).

Usage of the four CGN codons in human genes, despite the hypermutability of CpG dinucleotides and existence of AGR alternative codons, suggests the possibility of negative selective pressure at these sites. Arginine codon usage might therefore be a metric of interest for evaluation of disease genes. We hypothesized that genes rich in CGN codons are intolerant to variation and result in a greater burden of human disease, particularly for dominant phenotypes where one mutation can produce a phenotype.

## MATERIALS AND METHODS

### ClinVar SNV analysis

Unique ‘Pathogenic’ and ‘Benign’ nonsynonymous SNV entries were extracted from ClinVar variant summary data (accessed April 18, 2019). Using only entries in three letter amino acid code format (e.g. p.Arg330Met; 82.8% of all unique SNV entries), the number of amino acid substitutions and nucleotide changes corresponding to arginine substitutions were tallied. One entry of selenocysteine was omitted. Scripts used for these and the following analyses are available in **Supplementary Materials**.

### Pathogenic arginine codon usage analysis

AGR and CGN arginine codons were tallied for each Ensembl coding sequence transcript (GRCh38, release 96) (*5*) labeled ‘canonical’ in the Genome Aggregation Database (gnomAD, v2.1.1) (*6*). Disease-associated genes were selected from the Simons Foundation Autism Research Initiative (SFARI; https://www.sfari.org/resource/sfari-gene, accessed January 31, 2018), human homologs of essential genes found in a Drosophila melanogaster X chromosome screen (*7*), and the Online Mendelian Inheritance of Man (OMIM; https://www.omim.org/downloads, accessed April 9, 2018). To avoid ambiguity, only OMIM genes exclusively associated with dominant or recessive diseases were used; those associated with both were excluded, while those associated with neither were used as controls.

### Expression analysis

Using the GRCh38 reference genome (release 96), guanine and cytosine (GC) content was calculated as the proportion of bases that were guanine or cytosine 5 kb up- and 5 kb downstream of the canonical transcript boundaries, which were downloaded from in Ensembl BioMart. Gene expression data was downloaded from the Genotype-Tissue Expression (GTEx) database (accessed April 16, 2019) (*8*).

### Statistics

CGN codon frequencies were compared between disease-associated genes and those lacking disease association in their respective databases using a two-sided, two-sample Welch’s t-test or analysis of variants (ANOVA) in combination with Tukey’s honestly significant difference (HSD) tests. Pearson correlation was used to test relationships between CGN codon frequency by gene and gene constraint metrics listed in gnomAD (v2.1.1), GC content, and gene expression levels.

## RESULTS

Of all arginine codons found within 18,321 canonical gene transcripts in the human genome, the two AGR arginine codons were identified most frequently (AGA: 21.50%, AGG: 20.81%). However, the total frequency of these two AGR codons (42.31%) was below the total frequency of the four CGN codons (57.69%). Among CGN codons, CGG was the most frequent (20.40%), followed by CGC (18.42%), CGA (10.94%), and CGU (7.94%) (**Supplementary Table 1**). These results suggest that, as a group, there is not a dramatic depletion of CGN arginine codons, despite their relatively high risk of mutation.

Considering the unexpected persistence of CGN codons, we next analyzed CGN codon frequency in the context of putative variant pathogenicity. The frequency of CGN codon usage was higher in genes associated with syndromic autism spectrum disorder (ASD) as listed by SFARI (n=126) relative to the CGN arginine codon frequency in genes not in any way associated with ASD (n=17,369; p=0.0059, two-sided Welch’s t-test, **Figure 1C**). This was interesting given the high rate of de novo mutation in the ASD gene set (*9–11*). Likewise, human genes that are homologous to genes found in a *Drosophila melanogaster* X chromosome screen for essential genes (n=249) (*7*) also showed higher CGN codon frequency compared to all other genes (n=18,072; p=0.0001, two-sided Welch’s t-test, **Figure 1D**) suggesting that essential genes in model organisms are homologous to a gene set enriched for “high CGN” genes in the human genome.

To expand our analysis to monogenic Mendelian disorders, we next extracted a list of genes from OMIM that were exclusively annotated as ‘dominant’ (n=1,061) or ‘recessive’ (n=1,847). A comparison of these two gene groups alongside control genes that matched neither classification (n=14,946) revealed a significant difference in CGN arginine codon frequency (p=3.06E-22, ANOVA; **Figure 1E**) with a trend toward decreasing CGN frequency moving from dominant to recessive to control genes (dominant versus recessive: p=0.0009, Tukey HSD; dominant versus controls: p=3.39E-08, Tukey HSD; recessive versus controls: p=1.06E-07, Tukey HSD). Additionally, among genes with ClinVar SNV entries, we found that the proportion of ‘Pathogenic’ SNVs resulting in arginine substitutions was higher for genes with CGN codon frequencies in the top 25^th^ percentile (n=841) compared to genes with CGN frequencies in the bottom 25^th^ percentile (n=855; p=0.0093, two-sided Welch two sample t-test). Yet, there was no statistically significant difference in the total number of pathogenic SNVs (including all amino acid changes) (p=0.3914) or the total number of arginine codons per gene (p=0.4703). Similarly, the proportion of ‘Benign’ SNVs affecting arginine was higher in genes from the top 25^th^ percentile of CGN frequency (n=734) relative to genes from the bottom 25^th^ percentile (n=714; p=0.0036), while the total number of SNVs and total arginine codons did not differ (p=0.2446 and p=0.4022, respectively). However, the top ‘Benign’ genes showed a significantly lower proportion of total arginine SNVs than the top ‘Pathogenic’ SNVs (p<2.2E-16).

These results indicated that genes associated with genetic disorders tend to have higher CGN arginine codon frequencies. Further, our analysis of OMIM classifications, which revealed progressive differences between dominant and recessive genes, suggested that CGN codon frequency might inform how severe the effect of arginine substitutions might be, with a greater likelihood of impacting the phenotype of heterozygous individuals. To this end, we compared CGN codon frequency with different gnomAD constraint metrics. There was a modest, yet significant, positive correlation between CGN codon frequency and the probability of loss-of-function (pLI) score (r=0.099, p<2.2E-16, Pearson correlation; **Figure 1F**), which ranges from 0, very tolerant, to 1, not tolerant. This trend was stronger in a comparison of CGN frequency with observed-over-expected (o/e) ratios of loss-of-function (LoF) and missense mutations for each gene, showing a significant negative correlation in both instances (LoF: r=−0.128, p<2.2E-16, Pearson correlation, **Figure 1G**; missense: r=−0.163, p<2.2E-16, Pearson correlation, **Figure 1H**).

These data suggested that high CGN codon usage could point to regions under negative selective pressure. Yet, the mutability of CGN arginine residues might be balanced with higher GC content, which can impact gene expression (*3*). We therefore speculated that higher expressed genes in regions of high GC content might contribute to associations between CGN arginine codon frequency and human disease. We found a positive correlation between CGN codon frequency and the proportion of bases 5 kb up- and 5 kb downstream of the canonical transcript ends that were guanine or cytosine (r=0.537, p < 2.2E-16, Pearson correlation; **Figure 1I**). We next compared CGN codon frequency with the median transcripts per million (TPM) for each tissue in GTEx. With the exceptions of liver, minor salivary gland, stomach, testis, and whole blood, gene expression levels were positively correlated (Pearson) with CGN codon frequency in the 48 remaining tissues at a Bonferroni corrected significance threshold (p < 9.4E-4; **Figures 1J-K**).

Finally, we examined arginine codon usage in the context of a family of human disease genes, namely actin loci. We note that there are six actin paralogs in the human genome (*ACTG2*, *ACTA1*, *ACTA2*, *ACTC1*, *ACTB*, and *ACTG1*). Interestingly, all six human actin genes are Mendelian disease genes, and arginine substitutions at CGN codons have been identified as pathogenic alleles for all six genes (*ACTG2*: visceral myopathy, MIM#155310; *ACTA1*: nemaline myopathy, MIM#161800; *ACTA2:* familial thoracic aortic aneurysm, MIM#611788; *ACTC1*: cardiomyopathy and left ventricular noncompaction, MIM#613424; *ACTB*: Baraitser-Winter syndrome 1, MIM#243310; *ACTG1*: Baraitser-Winter syndrome 2, MIM#614583). Every actin paralog encodes 18 arginine residues that are highly conserved (**Figure 2A**) and encoded either by a CGN (**Figure 2**, red) or AGR trinucleotide (**Figure 2**, blue). Given the severity of the diseases phenotypes, we used the presence of missense substitution in the gnomAD database as a proxy for benign variation in each of the actin genes (**Figure 2A**). Interestingly, these benign variants all occurred in CGN codons rather than AGR codons (**Figure 2A**). We next compared these arginine residues across the actin paralogs to known pathogenic variants from ClinVar and discovered that all pathogenic alleles also occurred at CGN codons (**Figure 2B**). This near exclusive occurrence of both benign and pathogenic arginine codon variation at CGN codons can likely be attributed to their hypermutability relative to AGR codons; yet, distinct positions exhibited conserved CGN codon usage despite this hypermutability and were consistently mutated in multiple human diseases (e.g. arginine 12 and 14). In summary, the analysis of arginine codon usage could identify key hotspots for pathogenic alleles across a family of related human disease genes.

**Figure 2.**
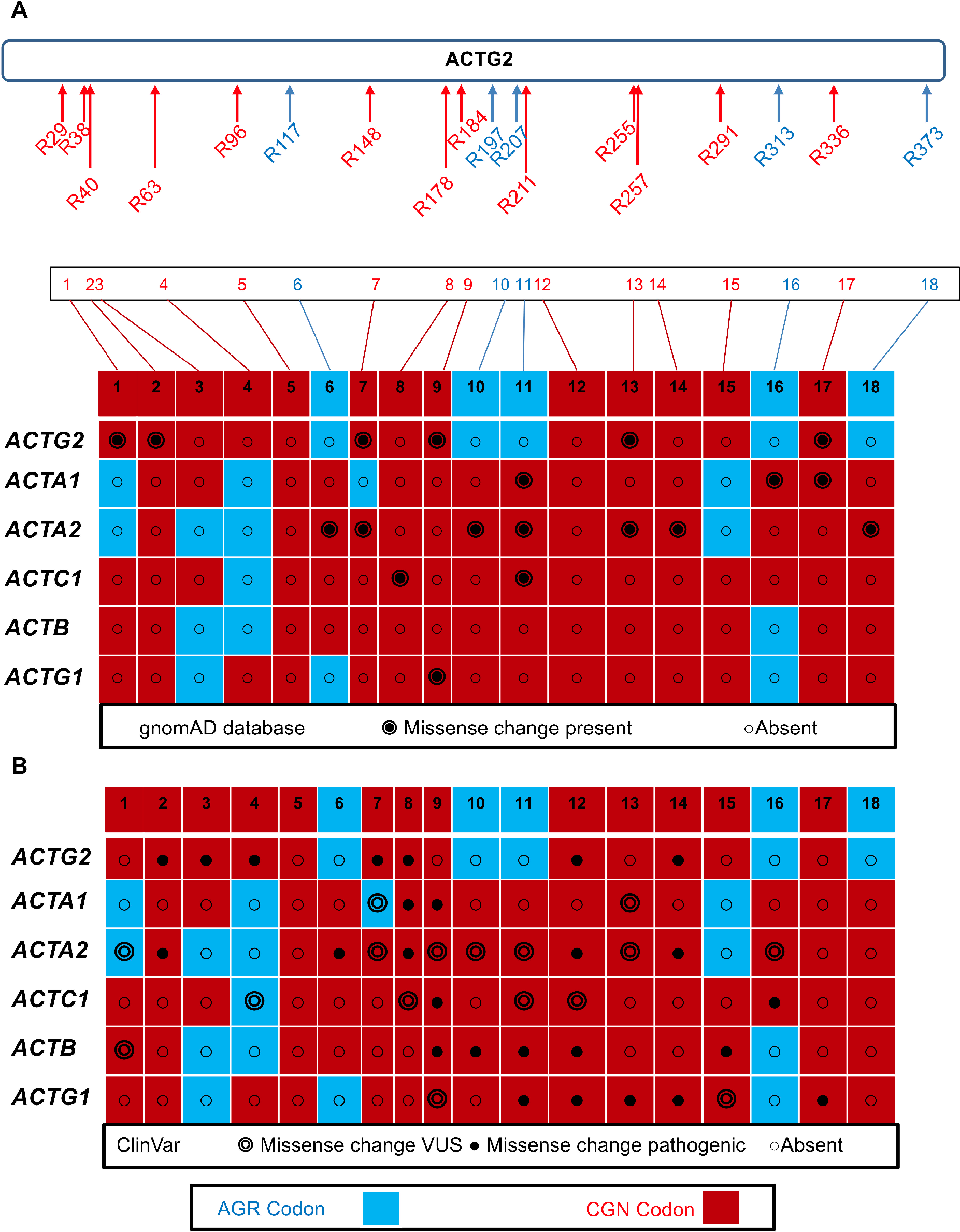
CGN arginine codon usage and variation across actin gene family. CGN codons are shown in red font or shading, AGR codons in blue. (**A**) Eighteen conserved actin residues in the *ACTG2* gene are shown at top with numbering corresponding to the ACTG2 protein. Each numbered arginine is then linked to a table of the actin paralogs. Solid black circle-dot marks indicate the presence of a missense change in the gnomAD database, a proxy for benign alleles. (**B**) Clinvar variants corresponding to the same arginine positions in A are shown for all homologs. Empty circle-dot symbols indicate variants of uncertain significance (VUS), while solid dots indicate pathogenic or likely pathogenic missense changes. Arginines 8, 9, 12, 13, and 14 are notable for being encoded by CGN codons in all paralogs and having pathogenic alleles across multiple human disease phenotypes.

## DISCUSSION

We analyzed coding regions of the human genome focused on arginine codons and found that genes with high proportions of CGN arginine codons compared to AGR were enriched among known disease genes and correlated with measures of mutation intolerance, indicating that genes with CGN codon preferences are more likely to underlie single gene disorders. Our results suggest that some genes have retained CGN codons that encode arginine despite their hypermutability and the often deleterious results on protein function that come from C>T transitions at those sites. As the only amino acid encoded by CpGs at the 1^st^ and 2^nd^ position, studying arginine codon usage provided us with some unique insights into these hot spots for mutation in disease. We speculate that the CGN codons have been anthropologically retained for reasons related to GC content and gene expression, consistent with our observation that genes with higher expression in brain and heart have biased CGN codon usage. Our results provide the possibility of simple gene-level predictions, but by pairing our analysis across highly identical paralogs (such as actin genes) we can see patterns of hypermutable arginine positions associated with disease.

We relied on large public databases, such as ClinVar and gnomAD, for our analyses, which on the one hand allowed us to take a broad, agnostic approach to the study of arginine codon usage; on the other hand, this meant that some of the limitations of these databases also extend to our study. Identifying SNVs as ‘Pathogenic’, for example, can include the use of damage prediction algorithms, which, depending on their heuristic approach, might more readily identify arginine substitutions as pathogenic compared to other amino acid changes and thus skew the results collected in ClinVar. Constraint metrics provided by gnomAD are limited by the populations included in its database, such that some genes might be more tolerant to variation than indicated simply because that variation is not seen in the sampled populations.

An outstanding question from our analyses is why such mutation-prone CGN arginine codons should persist in the human genome. One speculation is that high GC content might protect against deamination of methylated CpG sites by reducing the rate of DNA melting, thereby decreasing the amount of time spent in a single stranded state during which CpG sites are more vulnerable to deamination (*12*). Similarly, GC-biased gene conversion could maintain or increase the GC content, and thus CGN codon usage, in certain genomic regions (*13*). The positive correlation between CGN codon frequency and GC content observed in our study lends plausibility to both theories.

Classification of genetic variants with regard to their impact on human disease remains a major diagnostic challenge. Here we show that genes with a high frequency of CGN codons are enriched for human monogenic disease loci. Our observations of the relationship between human disease and CGN codon usage could be useful in more broadly predicting pathogenicity at the gene and variant level in gene discovery. Moreover, as these sites are highly likely to undergo mutations recurrently in the human population over time, the CGN codons are sites of highly recurrent pathogenic variants. Predicting the most likely mutation sites in disease could prove particularly useful in flagging specific recurrent alleles for oligonucleotide therapy.

## ACKNOWLEDGEMENTS

The authors would like to thank the Genome Aggregation Database (gnomAD) and the groups that provided exome and genome variant data to this resource. A full list of contributing groups can be found at https://gnomad.broadinstitute.org/about

## REFERENCES

1. T. Smith et al., Extensive Variation in the Mutation Rate Between and Within Human Genes Associated with Mendelian Disease. Hum Mutat 37, 488–494 (2016).

2. Z. Gao et al., Overlooked roles of DNA damage and maternal age in generating human germline mutations. Proc Natl Acad Sci U S A 116, 9491–9500 (2019).

3. S. Karlin, J. Mrazek, What drives codon choices in human genes? J Mol Biol 262, 459–472 (1996).

4. M. J. Landrum et al., ClinVar: public archive of interpretations of clinically relevant variants. Nucleic Acids Res 44, D862–868 (2016).

5. D. R. Zerbino et al., Ensembl 2018. Nucleic Acids Res 46, D754–d761 (2018).

6. M. Lek et al., Analysis of protein-coding genetic variation in 60,706 humans. Nature 536, 285–291 (2016).

7. S. Yamamoto et al., A drosophila genetic resource of mutants to study mechanisms underlying human genetic diseases. Cell 159, 200–214 (2014).

8. G. Consortium, The Genotype-Tissue Expression (GTEx) project. Nat Genet 45, 580–585 (2013).

9. I. Iossifov et al., The contribution of de novo coding mutations to autism spectrum disorder. Nature 515, 216–221 (2014).

10. T. N. Turner et al., Genomic Patterns of De Novo Mutation in Simplex Autism. Cell 171, 710–722.e712 (2017).

11. S. J. Sanders et al., De novo mutations revealed by whole-exome sequencing are strongly associated with autism. Nature 485, 237–241 (2012).

12. K. J. Fryxell, W. J. Moon, CpG mutation rates in the human genome are highly dependent on local GC content. Mol Biol Evol 22, 650–658 (2005).

13. F. Pouyet, D. Mouchiroud, L. Duret, M. Semon, Recombination, meiotic expression and human codon usage. Elife 6, (2017).

