## Supplementary material for "Biases in arginine codon usage correlate with genetic disease risk": step2_countCodons_fromFASTAformat_ensembl.py

```
import textwrap
import csv

# list of 'canonical' EMSEMBL identifiers that overlap with .fa filepath
canonicalIDs = [line.rstrip('\n') for line in \
open('ensemblIDs_overlap_Seq-gnomAD.txt')]

# list of ENSEMBL identifiers from .fa file
IDs = list()

# list of codons
codons = ["CGA", "CGC", "CGG", "CGT", "AGA", "AGG", "CTA", "CTC", "CTG", "CTT",
  "TTA", "TTG", "TCA", "TCC", "TCG", "TCT", "AGC", "AGT", "ACA", "ACC", "ACG",
  "ACT", "CCA", "CCC", "CCG", "CCT", "GCA", "GCC", "GCG", "GCT", "GGA", "GGC",
  "GGG", "GGT", "GTA", "GTC", "GTG", "GTT", "AAA", "AAG", "AAC", "AAT", "CAA",
  "CAG", "CAC", "CAT", "GAA", "GAG", "GAC", "GAT", "TAC", "TAT", "TGC", "TGT",
  "TTC", "TTT", "ATA", "ATC", "ATT", "ATG", "TGG", "TAA", "TAG", "TGA"]

# keep track of codon counts
counts = list()

# temporary sequence storage
seq = list()

# keep track if the current sequence is canonical
isCanonical = False

# Read CDS line by line, keep count as we go
#with open('test.fa') as fp:
with open('Homo_sapiens.GRCh38.cds.all.fa') as fp:
    for cnt, line in enumerate(fp):
        if line.startswith(">"):
            # get ensembl ID
            id = line.split(" ")[0].split(".")[0][1:]

            # only process ensembl IDs that are canonical
            if any(id in s for s in canonicalIDs):

                # count the number of codons for the last sequence
                # then store them in the counts list, except for the first line
                if len(IDs) > 0:
                    tempCounts = [IDs[-1]]
                    for c in codons:
                        tempCounts.append(seq.count(c))
                    counts.append(tempCounts)

                # set temporary sequence storage back to zero to count the
                # number of codons for the next sequence
                seq = list()

                # update canonical id tracker
                isCanonical = True

                # add new CDS ID to list of identifiers
                IDs.append(id)

            else:
                isCanonical = False

        # save codons for canonical genes
        if isCanonical and not line.startswith(">"):
            # separate nucleotide sequence into codons
            # and add to current sequence
            seq.extend(textwrap.wrap(line, 3))

# store counts for the last sequence in the file
tempCounts = [IDs[-1]]
for c in codons:
    tempCounts.append(seq.count(c))
counts.append(tempCounts)

# write all codon counts to file, first line with codons as header
codons.insert(0, "ENSEMBL_ID")    # add column header for IDs
counts.insert(0, codons)      # make codon info first line of output file
#with open('output.test_n2.csv', 'w') as out:
with open('output.ensembl-canonical.allCodonCounts.csv', 'w') as out:
    wr = csv.writer(out)
    wr.writerows(counts)
```
