## Supplementary material for "Biases in arginine codon usage correlate with genetic disease risk": step3_getArgCodonCounts_fromAllCodonCounts.R

```
###
### Arginine codon bias analysis with ensembl sequences
###

setwd("~/Documents/BCM Graduate School/Arginine Codon Usage/")

### load all codon counts
all <- read.csv("./ENSEMBL/output.ensembl-canonical.allCodonCounts.csv",
  header=T, stringsAsFactors = F)

### arginine codons
argCodons <- c("AGA", "AGG", "CGA", "CGC", "CGG", "CGT")

### calculate codon sums for each ensembl transcript
all$totalCodons <- rowSums(all[,2:ncol(all)])
all$totalArg <- rowSums(all[,argCodons])

### make arginine specific table
arg <- all[,c("ENSEMBL_ID", argCodons, "totalCodons", "totalArg")]

### calculate CG* and AG* codon frequency
arg$freqAG <- rowSums(arg[,c("AGA","AGG")]) / arg$totalArg
arg$freqCG <- rowSums(arg[,c("CGA","CGC","CGG","CGT")]) / arg$totalArg

### load gnomAD data
gnomad <- read.table("gnomAD/gnomad.v2.1.1.lof_metrics.by_transcript.txt",
  header=T, stringsAsFactors = F, sep="\t")
gnomad <- gnomad[gnomad$transcript %in% arg$ENSEMBL_ID,]
gnomad <- gnomad[order(match(gnomad$transcript, arg$ENSEMBL_ID)),]

### add information from gnomad data to arg count table
arg$gene <- gnomad$gene
arg$gene_ID <- gnomad$gene_id
arg <- arg[,c("ENSEMBL_ID", "gene_ID", "gene", colnames(arg)[2:(ncol(arg)-2)])]
arg$pLI.score <- gnomad$pLI
arg$oe_lof <- gnomad$oe_lof
arg$oe_mis <- gnomad$oe_mis

### for each arg codon, calculate which percentage of total arg codons
### it makes up
100 * (sum(arg$AGA)/sum(arg$totalArg))
100 * (sum(arg$AGG)/sum(arg$totalArg))
100 * (sum(arg$CGA)/sum(arg$totalArg))
100 * (sum(arg$CGC)/sum(arg$totalArg))
100 * (sum(arg$CGG)/sum(arg$totalArg))
100 * (sum(arg$CGT)/sum(arg$totalArg))

### save arg codon table
write.csv(arg, "ENSEMBL/argCodonCounts_ensembl_withGeneSymbols_withMetrics.csv",
  row.names=F, quote=F)
```
