## Supplementary material for "Biases in arginine codon usage correlate with genetic disease risk": step4_diseaseGenes_vs_Background.R

```
###
### Comparison with disease genes.
###

setwd("~/Documents/BCM Graduate School/Arginine Codon Usage/")

### load arg codon counts data
arg <- read.csv("ENSEMBL/argCodonCounts_ensembl_withGeneSymbols_withMetrics.csv",
  header=T, stringsAsFactors = F)

### SFARI
sfari <- read.csv("CCDS/genes_SFARI.csv", header=T)
sfariCGfreq <- arg$freqCG[arg$gene %in% unique(sfari$gene.symbol[sfari$syndromic == 1])]
non_sfariCGfreq <- arg$freqCG[!(arg$gene %in% unique(sfari$gene.symbol))]
asdp <- t.test(sfariCGfreq, non_sfariCGfreq)$p.value
asdtemp <- data.frame(freqCG=c(sfariCGfreq, non_sfariCGfreq),
                      group=factor(c(rep("SFARI", length(sfariCGfreq)), 
                                     rep("controls", length(non_sfariCGfreq))), 
                                   levels=c("SFARI", "controls")))

png("OMIM/CGfreq_bySyndromicSFARI.png",
  width=1450, height=1250, res=300)
par(mar=c(5.1,4.1,4.1,2.1))
boxplot(freqCG ~ group, data=asdtemp, ylim=c(0,1), 
        ylab="CGN codon frequency",xaxt="n", yaxt='n', xlab='',
        main="syndromic SFARI genes", cex.main=1.5, cex.lab=1.4)
axis(side=1, at=1:2, c(paste0("SFARI\n(n=", length(sfariCGfreq),")"), 
                       paste0("controls\n(n=", length(non_sfariCGfreq), ")")), 
     tick=FALSE, cex.axis=1.4, line=1)
axis(side=2, at=seq(0,1,0.25), labels=c('0.0', '', '0.5', '', '1.0'), 
     tick=T, cex.axis=1.4)
stripchart(freqCG ~ group, data=asdtemp, method='jitter', jitter=0.3, pch=c("."), 
           add=T, vertical=T, cex=1.75, col="black")
dev.off()

t.test(sfariCGfreq, non_sfariCGfreq)

### X-screen
x <- read.csv("CCDS/genes_Xscreen_fromPublication.csv", header=T)
xCGfreq <- arg$freqCG[arg$gene %in% unique(x$HumanGeneSymbol)]
non_xCGfreq <- arg$freqCG[!(arg$gene %in% unique(x$HumanGeneSymbol))]
xtemp <- data.frame(cgfreq=c(xCGfreq, non_xCGfreq),
  group=factor(c(rep("X-screen", length(xCGfreq)),
  rep("Non-X-screen", length(non_xCGfreq))), 
  levels=c("X-screen", "Non-X-screen")))

png("OMIM/CGfreq_byXscreen.png",
  width=1450, height=1250, res=300)
par(mar=c(5.1,4.1,4.1,2.1))
boxplot(cgfreq ~ group, data=xtemp, xaxt='n', ylab="CGN codon frequency",
  main="X-screen genes", cex.main=1.5, cex.lab=1.3, ylim=c(0,1), yaxt='n')
axis(side = 1,at = 1:2, c(paste0("X-screen\n(n=", length(xCGfreq), ")"), 
  paste0("controls\n(n=", length(non_xCGfreq), ")")),
  tick = FALSE, cex.axis=1.5, line=1.2)
axis(side=2, at=seq(0,1,0.25), labels=c('0.0', '', '0.5', '', '1.0'), 
     tick=T, cex.axis=1.4)
stripchart(cgfreq ~ group, data=xtemp, method='jitter', jitter=0.3, pch=c("."), 
           add=T, vertical=T, cex=1.75, col="black")
dev.off()

t.test(xCGfreq, non_xCGfreq)

### OMIM
omim <- read.xlsx("OMIM/genemap2.xlsx", sheetName="genemap2", 
                  startRow=4, endRow=16607, header=T)

dom_genes <- unique(unlist(strsplit(as.vector(omim$Gene.Symbol[grep(pattern="dominant", 
                                                                    x=omim$Phenotypes)]), ", ")))
rec_genes <- unique(unlist(strsplit(as.vector(omim$Gene.Symbol[grep(pattern="recessive", 
                                                                    x=omim$Phenotypes)]), 
                                    ", ")))
both <- intersect(dom_genes, rec_genes)

dom_exclusive <- dom_genes[!(dom_genes %in% both)]
rec_exclusive <- rec_genes[!(rec_genes %in% both)]

###
temp <- data.frame(gene = c(arg$gene[arg$gene %in% dom_exclusive],
                            arg$gene[arg$gene %in% rec_exclusive],
                            arg$gene[!(arg$gene %in% dom_exclusive) &
                                       !(arg$gene %in% rec_exclusive) &
                                       !(arg$gene %in% both)]),
                   freqCG = c(arg$freqCG[arg$gene %in% dom_exclusive],
                              arg$freqCG[arg$gene %in% rec_exclusive],
                              arg$freqCG[!(arg$gene %in% dom_exclusive) &
                                           !(arg$gene %in% rec_exclusive) &
                                           !(arg$gene %in% both)]),
                   group = c(rep("dominant", nrow(arg[arg$gene %in% dom_exclusive,])), 
                             rep("recessive", nrow(arg[arg$gene %in% rec_exclusive,])), 
                             rep("neither", nrow(arg[!(arg$gene %in% dom_exclusive) &
                                                       !(arg$gene %in% rec_exclusive) &
                                                       !(arg$gene %in% both),]))))
temp$group <- factor(temp$group, levels=c('dominant', 'recessive', 'neither'))

fit <- aov(freqCG ~ group, data=temp)
output <- TukeyHSD(fit)$group


png("ENSEMBL/CGfreq_byModeOfInheritanceOMIM_v1.png",
    width=1450, height=1250, res=300)
boxplot(freqCG ~ group, data=temp, 
        main="OMIM genetic disease", 
        ylab="CGN codon frequency",
        xaxt='n', xlab='', outcol="white", cex.main=1.5, cex.lab=1.3)
axis(side = 1,at = 1:3, c(paste0("dominant\n(n=", table(temp$group)["dominant"], ")"), 
                          paste0("recessive\n(n=", table(temp$group)["recessive"], ")"),
                          paste0("neither\n(n=", table(temp$group)["neither"], ")")),
     tick = FALSE, cex.axis=1.2, line=1)
stripchart(freqCG ~ group, data=temp, add=T, vertical=T,
           pch=".", col="black", method="jitter", jitter=0.3)
dev.off()
```
