## Supplementary material for "Biases in arginine codon usage correlate with genetic disease risk": step5_clinvarPathoBenign_TopVSBottomCGfreq.R

```
### Pathogenic variants
clinvar <- read.table(paste0("~/Documents/BCM Graduate School/",
  "Arginine Codon Usage/ClinVar/variant_summary_onlyPathogenicSNVs.txt"),
  header=T, sep="\t", stringsAsFactors=F)
clinvar <- unique(clinvar)

arg <- read.csv(paste0("~/Documents/BCM Graduate School/",
  "Arginine Codon Usage/ENSEMBL/argCodonCounts_ensembl_withGeneSymbols",
  "_withMetrics.csv"), stringsAsFactors=F)

arg$numPathoTotalSNVs <- rep(0, nrow(arg))
arg$numPathoArgSNVs <- rep(0, nrow(arg))
arg$propPathoArgSNVs <- rep(0, nrow(arg))

inter <- intersect(arg$gene, clinvar$GeneSymbol)
for(i in inter){
  temp <- clinvar[clinvar$GeneSymbol == i,]
  arg$numPathoTotalSNVs[arg$gene == i] <- nrow(temp)
  arg$numPathoArgSNVs[arg$gene == i] <- sum(grepl("(p.Arg", temp$Name, fixed=T))
  arg$propPathoArgSNVs[arg$gene == i] <-
    sum(grepl("(p.Arg", temp$Name, fixed=T)) / nrow(temp)
}


temp <- arg[arg$numPathoTotalSNVs != 0,]
temp <- temp[order(temp$freqCG, decreasing=T),]
#top <- temp[1:840,]
#bottom <- temp[2522:3361,]
top <- temp[temp$freqCG >= quantile(temp$freqCG, probs=0.75, na.rm=T),]
bottom <- temp[temp$freqCG <= quantile(temp$freqCG, probs=0.25, na.rm=T),]

t.test(top$numPathoTotalSNVs, bottom$numPathoTotalSNVs)
t.test(top$numPathoArgSNVs/top$numPathoTotalSNVs,
  bottom$numPathoArgSNVs/bottom$numPathoTotalSNVs)
t.test(top$totalArg, bottom$totalArg)


### Benign variants
benign <- read.table(paste0("~/Documents/BCM Graduate School/",
  "Arginine Codon Usage/ClinVar/variant_summary_onlyBenignSNVs.txt"),
  header=T, sep="\t", stringsAsFactors=F)
benign <- unique(benign)

arg$numBenignTotalSNVs <- rep(0, nrow(arg))
arg$numBenignArgSNVs <- rep(0, nrow(arg))
arg$propBenignArgSNVs <- rep(0, nrow(arg))

inter <- intersect(arg$gene, benign$GeneSymbol)
for(i in inter){
  temp <- benign[benign$GeneSymbol == i,]
  arg$numBenignTotalSNVs[arg$gene == i] <- nrow(temp)
  arg$numBenignArgSNVs[arg$gene == i] <- sum(grepl("(p.Arg", temp$Name, fixed=T))
  arg$propBenignArgSNVs[arg$gene == i] <-
    sum(grepl("(p.Arg", temp$Name, fixed=T)) / nrow(temp)
}

temp <- arg[arg$numBenignTotalSNVs != 0,]
temp <- temp[order(temp$freqCG, decreasing=T),]
#top <- temp[1:713,]
#bottom <- temp[2141:2853,]
top <- temp[temp$freqCG >= quantile(temp$freqCG, probs=0.75, na.rm=T),]
bottom <- temp[temp$freqCG <= quantile(temp$freqCG, probs=0.25, na.rm=T),]

t.test(top$numBenignTotalSNVs, bottom$numBenignTotalSNVs)
t.test(top$numBenignArgSNVs/top$numBenignTotalSNVs,
  bottom$numBenignArgSNVs/bottom$numBenignTotalSNVs)
t.test(top$totalArg, bottom$totalArg)

colnames(arg)[(ncol(arg)-5):ncol(arg)] <-
  paste0("clinvar.", colnames(arg)[(ncol(arg)-5):ncol(arg)])

write.csv(arg, paste0("Documents/BCM Graduate School/",
  "Arginine Codon Usage/ENSEMBL/argCodonCounts_ensembl_withGeneSymbols",
  "_withMetrics_withClinVar.csv"), quote=F, row.names=F)

temp <- arg[arg$clinvar.numPathoTotalSNVs != 0,]
temp <- temp[order(temp$freqCG, decreasing=T),]
#toppatho <- temp[1:840,]
toppatho <- temp[temp$freqCG >= quantile(temp$freqCG, probs=0.75, na.rm=T),]

temp <- arg[arg$clinvar.numBenignTotalSNVs != 0,]
temp <- temp[order(temp$freqCG, decreasing=T),]
#topbenign <- temp[1:713,]
topbenign <- temp[temp$freqCG >= quantile(temp$freqCG, probs=0.75, na.rm=T),]

t.test(toppatho$clinvar.propPathoArgSNVs, topbenign$clinvar.propBenignArgSNVs)
```
