## Supplementary material for "Biases in arginine codon usage correlate with genetic disease risk": step6_CGfreq_byDamagingMetrics.R

```
###
### Compare CGN codon frequency to 'damaging' metrics
###

setwd("~/Documents/BCM Graduate School/Arginine Codon Usage/")

### load arg codon counts data
arg <- read.csv("ENSEMBL/argCodonCounts_ensembl_withGeneSymbols_withMetrics.csv",
  header=T, stringsAsFactors = F)

### CGN codon frequency versus pLI score
cor.test(arg$pLI.score, arg$freqCG)

mpli <- c(mean(arg$pLI.score[arg$freqCG >= 0 & arg$freqCG <= 0.05], na.rm=T),
  mean(arg$pLI.score[arg$freqCG > 0.05 & arg$freqCG <= 0.1], na.rm=T),
  mean(arg$pLI.score[arg$freqCG > 0.1 & arg$freqCG <= 0.15], na.rm=T),
  mean(arg$pLI.score[arg$freqCG > 0.15 & arg$freqCG <= 0.2], na.rm=T),
  mean(arg$pLI.score[arg$freqCG > 0.2 & arg$freqCG <= 0.25], na.rm=T),
  mean(arg$pLI.score[arg$freqCG > 0.25 & arg$freqCG <= 0.3], na.rm=T),
  mean(arg$pLI.score[arg$freqCG > 0.3 & arg$freqCG <= 0.35], na.rm=T),
  mean(arg$pLI.score[arg$freqCG > 0.35 & arg$freqCG <= 0.4], na.rm=T),
  mean(arg$pLI.score[arg$freqCG > 0.4 & arg$freqCG <= 0.45], na.rm=T),
  mean(arg$pLI.score[arg$freqCG > 0.45 & arg$freqCG <= 0.5], na.rm=T),
  mean(arg$pLI.score[arg$freqCG > 0.5 & arg$freqCG <= 0.55], na.rm=T),
  mean(arg$pLI.score[arg$freqCG > 0.55 & arg$freqCG <= 0.6], na.rm=T),
  mean(arg$pLI.score[arg$freqCG > 0.6 & arg$freqCG <= 0.65], na.rm=T),
  mean(arg$pLI.score[arg$freqCG > 0.65 & arg$freqCG <= 0.7], na.rm=T),
  mean(arg$pLI.score[arg$freqCG > 0.7 & arg$freqCG <= 0.75], na.rm=T),
  mean(arg$pLI.score[arg$freqCG > 0.75 & arg$freqCG <= 0.8], na.rm=T),
  mean(arg$pLI.score[arg$freqCG > 0.8 & arg$freqCG <= 0.85], na.rm=T),
  mean(arg$pLI.score[arg$freqCG > 0.85 & arg$freqCG <= 0.9], na.rm=T),
  mean(arg$pLI.score[arg$freqCG > 0.9 & arg$freqCG <= 0.95], na.rm=T),
  mean(arg$pLI.score[arg$freqCG > 0.95 & arg$freqCG <= 1.00], na.rm=T))

png("ENSEMBL/CGfreq_byPLI.png", height=1200, width=1200, res=300)
par(mar=c(5.1,4.1,4.1,4.1))
hist(arg$freqCG, xlab="CGN codon frequency", ylab="count, canonical genes",
  main="CGN codon frequency\nvs mean pLI score")
par(new=T)
plot(seq(0.025,0.975,0.05), mpli, col="blue", xlim=c(0,1), pch=19, 
  xaxt='n', yaxt='n',ylab="",xlab="",bty='n', ylim=c(0.1,0.4))
axis(4, col="blue", col.ticks="blue", col.axis="blue")
mtext("mean pLI score", side=4, line=2.5, col="blue")
dev.off()


### CGN codon frequency versus observed/expected LoF ratio
cor.test(arg$oe_lof, arg$freqCG)

mlof <- c(mean(arg$oe_lof[arg$freqCG >= 0 & arg$freqCG <= 0.05], na.rm=T),
  mean(arg$oe_lof[arg$freqCG > 0.05 & arg$freqCG <= 0.1], na.rm=T),
  mean(arg$oe_lof[arg$freqCG > 0.1 & arg$freqCG <= 0.15], na.rm=T),
  mean(arg$oe_lof[arg$freqCG > 0.15 & arg$freqCG <= 0.2], na.rm=T),
  mean(arg$oe_lof[arg$freqCG > 0.2 & arg$freqCG <= 0.25], na.rm=T),
  mean(arg$oe_lof[arg$freqCG > 0.25 & arg$freqCG <= 0.3], na.rm=T),
  mean(arg$oe_lof[arg$freqCG > 0.3 & arg$freqCG <= 0.35], na.rm=T),
  mean(arg$oe_lof[arg$freqCG > 0.35 & arg$freqCG <= 0.4], na.rm=T),
  mean(arg$oe_lof[arg$freqCG > 0.4 & arg$freqCG <= 0.45], na.rm=T),
  mean(arg$oe_lof[arg$freqCG > 0.45 & arg$freqCG <= 0.5], na.rm=T),
  mean(arg$oe_lof[arg$freqCG > 0.5 & arg$freqCG <= 0.55], na.rm=T),
  mean(arg$oe_lof[arg$freqCG > 0.55 & arg$freqCG <= 0.6], na.rm=T),
  mean(arg$oe_lof[arg$freqCG > 0.6 & arg$freqCG <= 0.65], na.rm=T),
  mean(arg$oe_lof[arg$freqCG > 0.65 & arg$freqCG <= 0.7], na.rm=T),
  mean(arg$oe_lof[arg$freqCG > 0.7 & arg$freqCG <= 0.75], na.rm=T),
  mean(arg$oe_lof[arg$freqCG > 0.75 & arg$freqCG <= 0.8], na.rm=T),
  mean(arg$oe_lof[arg$freqCG > 0.8 & arg$freqCG <= 0.85], na.rm=T),
  mean(arg$oe_lof[arg$freqCG > 0.85 & arg$freqCG <= 0.9], na.rm=T),
  mean(arg$oe_lof[arg$freqCG > 0.9 & arg$freqCG <= 0.95], na.rm=T),
  mean(arg$oe_lof[arg$freqCG > 0.95 & arg$freqCG <= 1.00], na.rm=T))

png("ENSEMBL/CGfreq_byOE-LoF.png", height=1200, width=1200, res=300)
par(mar=c(5.1,4.1,4.1,4.1))
hist(arg$freqCG, xlab="CGN codon frequency", ylab="count, canonical genes",
  main="CGN codon frequency\nvs mean o/e LoF")
par(new=T)
plot(seq(0.025,0.975,0.05), mlof, col="blue", xlim=c(0,1), pch=19, 
  xaxt='n', yaxt='n',ylab="",xlab="",bty='n', ylim=c(0.4,1.0))
axis(4, col="blue", col.ticks="blue", col.axis="blue")
mtext("mean o/e LoF (gnomAD)", side=4, line=2.5, col="blue")
dev.off()

### CGN codon frequency versus observed/expected missense ratio
cor.test(arg$oe_mis, arg$freqCG)

mmis <- c(mean(arg$oe_mis[arg$freqCG >= 0 & arg$freqCG <= 0.05], na.rm=T),
  mean(arg$oe_mis[arg$freqCG > 0.05 & arg$freqCG <= 0.1], na.rm=T),
  mean(arg$oe_mis[arg$freqCG > 0.1 & arg$freqCG <= 0.15], na.rm=T),
  mean(arg$oe_mis[arg$freqCG > 0.15 & arg$freqCG <= 0.2], na.rm=T),
  mean(arg$oe_mis[arg$freqCG > 0.2 & arg$freqCG <= 0.25], na.rm=T),
  mean(arg$oe_mis[arg$freqCG > 0.25 & arg$freqCG <= 0.3], na.rm=T),
  mean(arg$oe_mis[arg$freqCG > 0.3 & arg$freqCG <= 0.35], na.rm=T),
  mean(arg$oe_mis[arg$freqCG > 0.35 & arg$freqCG <= 0.4], na.rm=T),
  mean(arg$oe_mis[arg$freqCG > 0.4 & arg$freqCG <= 0.45], na.rm=T),
  mean(arg$oe_mis[arg$freqCG > 0.45 & arg$freqCG <= 0.5], na.rm=T),
  mean(arg$oe_mis[arg$freqCG > 0.5 & arg$freqCG <= 0.55], na.rm=T),
  mean(arg$oe_mis[arg$freqCG > 0.55 & arg$freqCG <= 0.6], na.rm=T),
  mean(arg$oe_mis[arg$freqCG > 0.6 & arg$freqCG <= 0.65], na.rm=T),
  mean(arg$oe_mis[arg$freqCG > 0.65 & arg$freqCG <= 0.7], na.rm=T),
  mean(arg$oe_mis[arg$freqCG > 0.7 & arg$freqCG <= 0.75], na.rm=T),
  mean(arg$oe_mis[arg$freqCG > 0.75 & arg$freqCG <= 0.8], na.rm=T),
  mean(arg$oe_mis[arg$freqCG > 0.8 & arg$freqCG <= 0.85], na.rm=T),
  mean(arg$oe_mis[arg$freqCG > 0.85 & arg$freqCG <= 0.9], na.rm=T),
  mean(arg$oe_mis[arg$freqCG > 0.9 & arg$freqCG <= 0.95], na.rm=T),
  mean(arg$oe_mis[arg$freqCG > 0.95 & arg$freqCG <= 1.00], na.rm=T))

png("ENSEMBL/CGfreq_byOE-Missense.png", height=1200, width=1200, res=300)
par(mar=c(5.1,4.1,4.1,4.1))
hist(arg$freqCG, xlab="CGN codon frequency", ylab="count, canonical genes",
  main="CGN codon frequency\nvs mean o/e missense")
par(new=T)
plot(seq(0.025,0.975,0.05), mmis, col="blue", xlim=c(0,1), pch=19, 
  xaxt='n', yaxt='n',ylab="",xlab="",bty='n', ylim=c(0.4,1.0))
axis(4, col="blue", col.ticks="blue", col.axis="blue")
mtext("mean o/e missense (gnomAD)", side=4, line=2.5, col="blue")
dev.off()
```
