## Supplementary material for "Biases in arginine codon usage correlate with genetic disease risk": step1_getUpDownStreamTranscriptCoordinates.R

```
###
### Create .bed files with coordinates +/- 5kb of transcript starts
###

setwd("~/Documents/BCM Graduate School/Arginine Codon Usage/")

### load arg codon counts data
arg <- read.csv("ENSEMBL/argCodonCounts_ensembl_withGeneSymbols_withMetrics.csv",
  header=T, stringsAsFactors = F)

### load ensembl transcript coordinates
### file was downloaded from ENSEMBL biomart selecting the following:
### 1. Dataset: Human genes (GRCh38.p12)
### 2. Attributes: Gene stable ID, Transcript stable ID,
###      Transcript start (bp), Transcript end (bp),
###      Chromosome/scaffold name
coord <- read.csv("ENSEMBL/ensembl_transcript-coordinates.csv",
  header=T, stringsAsFactors = F)

### keep only those transcripts that overlap the ones in the arg table
coord <- coord[coord$Transcript.stable.ID %in% arg$ENSEMBL_ID,]
coord <- coord[order(match(coord$Transcript.stable.ID, arg$ENSEMBL_ID)),]

### 5 kb upstream
up <- as.data.frame(cbind(coord$Chromosome.scaffold.name,
  coord$Transcript.start..bp. - 5000, coord$Transcript.start..bp.,
  coord$Transcript.stable.ID))

### 5 kb downstream
down <- as.data.frame(cbind(coord$Chromosome.scaffold.name,
  coord$Transcript.end..bp., coord$Transcript.end..bp. + 5000,
  coord$Transcript.stable.ID))

### save output
write.table(up, "ENSEMBL/ensembl_upstreamOfTranscript_5k.bed",
  quote=F, sep="\t", row.names=F, col.names=F)
write.table(down, "ENSEMBL/ensembl_downstreamOfTranscript_5k.bed",
  quote=F, sep="\t", row.names=F, col.names=F)
```
