## Supplementary material for "Biases in arginine codon usage correlate with genetic disease risk": step3_CGfreq_byGCcontent_5kb-up-down.R

```
###
### CGN codon frequency versus GC content 5kb up- and downstream
### of transcripts.
###

setwd("~/Documents/BCM Graduate School/Arginine Codon Usage/")

### load arg codon counts data
arg <- read.csv("ENSEMBL/argCodonCounts_ensembl_withGeneSymbols_withMetrics.csv",
  header=T, stringsAsFactors = F)

### load up/downstream GC counts
up <- read.csv("ENSEMBL/upGC.csv", stringsAsFactors = F, header=F)
colnames(up) <- c("ENSEMBL_ID", "count")
down <- read.csv("ENSEMBL/downGC.csv", stringsAsFactors = F, header=F)
colnames(down) <- c("ENSEMBL_ID", "count")

### only assess transcripts with up AND down GC counts
both <- intersect(up$ENSEMBL_ID, down$ENSEMBL_ID)
up <- up[up$ENSEMBL_ID %in% both,]
down <- down[down$ENSEMBL_ID %in% both,]

### add GC count info to arg table
arg$GCcounts.5kb.up <- rep(NA, nrow(arg))
arg$GCcounts.5kb.down <- rep(NA, nrow(arg))
arg$GCcounts.5kb.up[arg$ENSEMBL_ID %in% both] <- up$count
arg$GCcounts.5kb.down[arg$ENSEMBL_ID %in% both] <- down$count

### summarize GC count info
arg$percGC.total <- rep(NA, nrow(arg))
arg$percGC.total[arg$ENSEMBL_ID %in% up$ENSEMBL_ID] <- 
  100 * ((arg$GCcounts.5kb.up[arg$ENSEMBL_ID %in% up$ENSEMBL_ID] +
     arg$GCcounts.5kb.down[arg$ENSEMBL_ID %in% down$ENSEMBL_ID])/10000)
cor.test(arg$percGC.total, arg$freqCG)

### plot GC content by CGN codon frequency
### CGN codon frequency versus pLI score
mgc <- c(mean(arg$percGC.total[arg$freqCG >= 0 & arg$freqCG <= 0.05], na.rm=T),
  mean(arg$percGC.total[arg$freqCG > 0.05 & arg$freqCG <= 0.1], na.rm=T),
  mean(arg$percGC.total[arg$freqCG > 0.1 & arg$freqCG <= 0.15], na.rm=T),
  mean(arg$percGC.total[arg$freqCG > 0.15 & arg$freqCG <= 0.2], na.rm=T),
  mean(arg$percGC.total[arg$freqCG > 0.2 & arg$freqCG <= 0.25], na.rm=T),
  mean(arg$percGC.total[arg$freqCG > 0.25 & arg$freqCG <= 0.3], na.rm=T),
  mean(arg$percGC.total[arg$freqCG > 0.3 & arg$freqCG <= 0.35], na.rm=T),
  mean(arg$percGC.total[arg$freqCG > 0.35 & arg$freqCG <= 0.4], na.rm=T),
  mean(arg$percGC.total[arg$freqCG > 0.4 & arg$freqCG <= 0.45], na.rm=T),
  mean(arg$percGC.total[arg$freqCG > 0.45 & arg$freqCG <= 0.5], na.rm=T),
  mean(arg$percGC.total[arg$freqCG > 0.5 & arg$freqCG <= 0.55], na.rm=T),
  mean(arg$percGC.total[arg$freqCG > 0.55 & arg$freqCG <= 0.6], na.rm=T),
  mean(arg$percGC.total[arg$freqCG > 0.6 & arg$freqCG <= 0.65], na.rm=T),
  mean(arg$percGC.total[arg$freqCG > 0.65 & arg$freqCG <= 0.7], na.rm=T),
  mean(arg$percGC.total[arg$freqCG > 0.7 & arg$freqCG <= 0.75], na.rm=T),
  mean(arg$percGC.total[arg$freqCG > 0.75 & arg$freqCG <= 0.8], na.rm=T),
  mean(arg$percGC.total[arg$freqCG > 0.8 & arg$freqCG <= 0.85], na.rm=T),
  mean(arg$percGC.total[arg$freqCG > 0.85 & arg$freqCG <= 0.9], na.rm=T),
  mean(arg$percGC.total[arg$freqCG > 0.9 & arg$freqCG <= 0.95], na.rm=T),
  mean(arg$percGC.total[arg$freqCG > 0.95 & arg$freqCG <= 1.00], na.rm=T))

png("ENSEMBL/CGfreq_byGC-content_up-down-5kb.png", 
  height=1200, width=1200, res=300)
par(mar=c(5.1,4.1,4.1,4.1))
hist(arg$freqCG, xlab="CGN codon frequency", ylab="count, canonical transcripts",
  main="CGN codon frequency\nvs mean percent GC content (+/- 5kb)")
par(new=T)
plot(seq(0.025,0.975,0.05), mgc, col="blue", xlim=c(0,1), pch=19, 
  xaxt='n', yaxt='n',ylab="",xlab="",bty='n', ylim=c(40,54))
axis(4, col="blue", col.ticks="blue", col.axis="blue")
mtext("mean percent GC", side=4, line=2.5, col="blue")
dev.off()
```
