## Supplementary material for "Biases in arginine codon usage correlate with genetic disease risk": step4_CGfreq_byGTExGeneExpression.R

```
###
### Compare CGN codon frequency to GTEx median TPM gene expression
###

setwd("~/Documents/BCM Graduate School/Arginine Codon Usage/")

### load arg codon counts data
arg <- read.csv("ENSEMBL/argCodonCounts_ensembl_withGeneSymbols_withMetrics.csv",
  header=T, stringsAsFactors = F)

### load GTEx data
gtex <- read.table("GTEx/GTEx_Analysis_2016-01-15_v7_RNASeQCv1.1.8_gene_median_tpm.gct",
  header=T, skip=2, sep="\t", stringsAsFactors = F)
gtex$gene_id <- unlist(strsplit(gtex$gene_id, "\\."))[c(T,F)]

### get overlapping gene IDs
inter <- intersect(arg$gene_ID, gtex$gene_id)

### reduce arg and gtex to overlapping gene IDs
arg <- arg[arg$gene_ID %in% inter,]
gtex <- gtex[gtex$gene_id %in% inter,]
gtex <- gtex[order(match(gtex$gene_id, arg$gene_ID)),]
gtex$freqCG <- arg$freqCG

### test correlation
p.values <- apply(gtex[,3:55], 2, function(x){cor.test(x, gtex$freqCG)$p.value})
table(p.values < (0.05/length(p.values)))
names(p.values)[!(p.values < (0.05/length(p.values)))]

### generate plot for each GTEx tissue
tissues <- colnames(gtex)[3:ncol(gtex)]
for(t in tissues){
  name <- gsub("\\.", " ", gsub("\\.\\.", " ", gsub("\\.\\.\\.", " ", t)))
  m <- c(mean(gtex[gtex$freqCG >= 0 & gtex$freqCG <= 0.05, t], na.rm=T),
    mean(gtex[gtex$freqCG > 0.05 & gtex$freqCG <= 0.1, t], na.rm=T),
    mean(gtex[gtex$freqCG > 0.1 & gtex$freqCG <= 0.15, t], na.rm=T),
    mean(gtex[gtex$freqCG > 0.15 & gtex$freqCG <= 0.2, t], na.rm=T),
    mean(gtex[gtex$freqCG > 0.2 & gtex$freqCG <= 0.25, t], na.rm=T),
    mean(gtex[gtex$freqCG > 0.25 & gtex$freqCG <= 0.3, t], na.rm=T),
    mean(gtex[gtex$freqCG > 0.3 & gtex$freqCG <= 0.35, t], na.rm=T),
    mean(gtex[gtex$freqCG > 0.35 & gtex$freqCG <= 0.4, t], na.rm=T),
    mean(gtex[gtex$freqCG > 0.4 & gtex$freqCG <= 0.45, t], na.rm=T),
    mean(gtex[gtex$freqCG > 0.45 & gtex$freqCG <= 0.5, t], na.rm=T),
    mean(gtex[gtex$freqCG > 0.5 & gtex$freqCG <= 0.55, t], na.rm=T),
    mean(gtex[gtex$freqCG > 0.55 & gtex$freqCG <= 0.6, t], na.rm=T),
    mean(gtex[gtex$freqCG > 0.6 & gtex$freqCG <= 0.65, t], na.rm=T),
    mean(gtex[gtex$freqCG > 0.65 & gtex$freqCG <= 0.7, t], na.rm=T),
    mean(gtex[gtex$freqCG > 0.7 & gtex$freqCG <= 0.75, t], na.rm=T),
    mean(gtex[gtex$freqCG > 0.75 & gtex$freqCG <= 0.8, t], na.rm=T),
    mean(gtex[gtex$freqCG > 0.8 & gtex$freqCG <= 0.85, t], na.rm=T),
    mean(gtex[gtex$freqCG > 0.85 & gtex$freqCG <= 0.9, t], na.rm=T),
    mean(gtex[gtex$freqCG > 0.9 & gtex$freqCG <= 0.95, t], na.rm=T),
    mean(gtex[gtex$freqCG > 0.95 & gtex$freqCG <= 1.00, t], na.rm=T))
  
  png(paste0("ENSEMBL/CGfreq_by_geneExpression_GTEx_", gsub(" ", "-", name),
    ".png"), height=1200, width=1200, res=300)
  par(mar=c(5.1,4.1,4.1,4.1))
  hist(gtex$freqCG, xlab="CGN codon frequency", ylab="count, canonical genes",
    main=paste0("CGN codon frequency\nvs ", name, " expression"))
  par(new=T)
  plot(seq(0.025,0.975,0.05), m, col="blue", xlim=c(0,1), pch=19, 
    xaxt='n', yaxt='n',ylab="",xlab="",bty='n')
  axis(4, col="blue", col.ticks="blue", col.axis="blue")
  mtext("mean expression", side=4, line=2.5, col="blue")
  dev.off()
}
```
